# A Protocol for Neuralized Murine Olfactory Organoids

**DOI:** 10.1101/2024.10.29.620938

**Authors:** Alp Ozgun, Priya Suman, Josée Coulombe, aSCENT-PD Investigators, Earl G. Brown, Julianna J. Tomlinson, John M. Woulfe, Michael G. Schlossmacher

## Abstract

Chronic olfactory dysfunction can be associated with parkinsonism, dementia, demyelinating disorders and schizophrenia. The olfactory epithelium (OE) represents an interface between the environment and the central nervous system. Mounting evidence implicates environmental factors in neurodegenerative disease processes, necessitating investigations into their interactions with the host’s genome. In Parkinson disease, hyposmia often precedes motor symptoms, raising the possibility that the OE could be involved in disease initiation. We previously demonstrated abundant α-synuclein expression in mammalian OE as well as aggregate formation in the olfactory nerve. Current *in vitro* models of OE are limited, relying primarily on post-mitotic cultures established from biopsies. To address this gap, we present a method for generating olfactory organoids of OE from adult mice. These organoids comprise neuronal and non-neuronal cell types, including sustentacular cells, thus encompassing structural elements of OE *in situ*. Expression of the olfactory sensory neuron marker OMP and Parkinson’s-linked α-synuclein was also detected in olfactory organoids, highlighting their potential usefulness to mechanistic research. We established OE organoids that were kept in culture for up to 3 weeks. In addition, we inoculated organoids with the neurotropic vesicular stomatitis virus to model infections. We conclude that this olfactory organoid model system offers a new platform for studying airborne environmental factors in their interactions with a genetically defined host; this, to study OE biology and enable the exploration of disease processes within olfactory tissue.

## Introduction

Studying the olfactory system has significant implications for unraveling the pathophysiology of various medical conditions and neurological diseases. The olfactory system serves as a unique window into neurological disorders due to its direct connection to the brain and its vulnerability to pathological changes. Importantly, olfactory dysfunction frequently manifests as an early and sometimes prodromal sign of numerous neurological disorders, including Alzheimer’s disease, Parkinson disease, multiple sclerosis, and schizophrenia. Moreover, the olfactory system is uniquely susceptible to environmental exposures due to its direct contact with airborne particles, including potential neurotoxicants, microbial pathogens and microplastics. This susceptibility is attributed to the location of the olfactory epithelium (OE) in the roof of the human nasal cavity and short axons traversing the base of the skull, thus potentially allowing environmental agents easier access to neural tissue.

Mounting evidence suggests that many neurodegenerative diseases, such as Alzheimer’s disease, Parkinson disease, and amyotrophic lateral sclerosis, are influenced by environmental factors, including pesticides^1,2^, air pollutants^3,4^ and infectious illnesses^5,6^. For example, a recent study identified an association between particular leisure activities, including golfing, hunting and gardening, and an elevated incidence of ALS^7^. This evidence underscores the significance of environmental exposures in the pathogenesis of neurodegenerative diseases and highlights the necessity for comprehensive investigations into diverse exposure factors within this framework. In the context of Parkinson disease, hyposmia often precedes motor symptoms by decades, identifying its potential as an early risk factor and implicating initiation mechanisms in the OE. Our prior research has shown that α-synuclein, which is associated with both heritable and sporadic forms of Parkinson’s, is abundantly expressed in the OE and contributes to antimicrobial defense mechanisms to protect against a nasally acquired viral infection^8^, thereby reinforcing the importance of investigating the OE in elucidating Parkinson disease pathophysiology.

The olfactory epithelium exhibits structural and phenotypic similarities to respiratory epithelial structures; it is characterized by the presence of sustentacular and basal cells expressing the markers expressed by respiratory epithelium cells, as illustrated in **Figure 1a**. However, a key distinction lies in the even more extensive neuralization of the OE. Our prior work utilized a full skull sectioning method for immunohistochemistry and immunofluorescence, facilitating the visualization of olfactory elements in their natural spatial relationship within intact murine skull sections^8^. Utilizing this method, we demonstrate, as shown in **Figure 1b**, the positioning of somata for mature and immature olfactory sensory neuron (OSN) within the OE. Mature dendritic knobs of OSNs carrying odor receptors are embedded in a thin layer of mucus and interface with air flow in the roof of the nasal cavity, while their axons penetrate the mucosal lamina propria, are collated into cranial nerve-I (CN-I) bundles and extend to glomeruli within the olfactory bulb. Sustentacular cells exhibit positive staining for CK18 (**Figure 1c**), while mature OSNs express olfactory marker protein (OMP) in their dendritic knobs and soma. Notably, beta-III tubulin (TUBB3) labels immature OSNs, with axons of CN-I displaying TUBB3-positivity and variable degrees of OMP expression. The latter observation corroborates previous reports^9^ indicating that immature OSNs lacking OMP can still establish neural connections and provide sensory input to the olfactory bulb.

**Figure 1:**
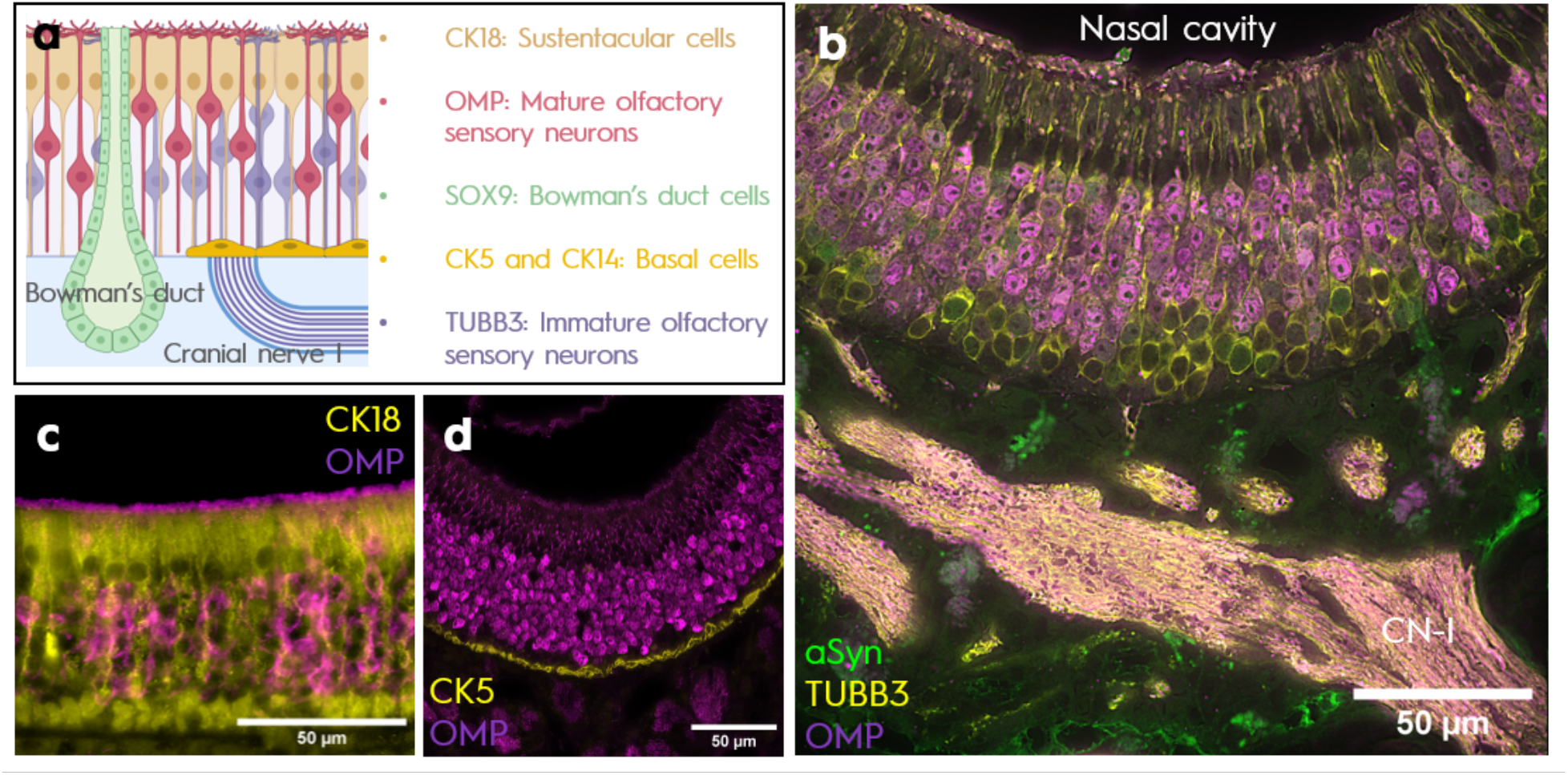
Structure of the murine olfactory epithelium. **a)** Schematic representation illustrating the cellular arrangement within the olfactory epithelium (OE), including a mucus-secreting Bowman’s gland and cranial nerve-I fibers. **b)** Intact olfactory epithelial section stained for immature (using anti-TUBB3) and mature (anti-OMP) olfactory sensory neurons, revealing the distribution of neuronal somata and composition of cranial nerve-I (CN-I) axons. Anti-α-synuclein staining highlights neuronal vs. non-neuronal expression. **c)** Staining for CK18 demonstrates the morphology of sustentacular cells. **d)** Staining for CK5 reveals the layer of horizontal basal cells. Images are taken from 6 month-old wild-type C57BL/6J mice.

Two distinct types of basal cells populate the OE: horizontal basal cells and globose basal cells. The latter are identified by their expression of the LGR5 surface marker and serve as a mitotic pool responsible for the routine turnover of other cell types within the OE. In contrast, horizontal basal cells, as depicted in **Figure 1d**, are a monolayer of dormant stem cells expressing CK5 and CK14 markers. In response to OE injury, these cells become activated and contribute to the replenishment of the globose basal cell pool, facilitating tissue regeneration. The *in vitro* expansion of these stem cells using a defined medium formulation supplemented with dual SMAD inhibitors has been previously reported and extensively characterized^10^.

The availability of *in vitro* models for mammalian OE are currently limited, predominantly relying on primary cultures derived from OE biopsies^11^. These cultures, however, are post-mitotic and lack the capacity for expansion, necessitating fresh biopsies for each experiment^12^. Historically, these cultures have been used to investigate the impact of air pollution^13^ and to characterize features of OSNs^13^. Another *in vitro* modeling approach involves differentiating OSNs from stem cells isolated from OE tissue collections via cell sorting^14^. More recent endeavors in modeling OE include air-liquid interface cultures, which have been utilized to explore the effects of pollution^15^ and any mechanistic links between SARS-CoV-2 infections and Alzheimer’s disease pathogenesis^16^. Nevertheless, to date, these cultures do not recapitulate neuronal phenotypes and are more suitable for investigating non-neuronal cell types.

Organoids are playing an increasingly important role in preclinical research, including studies into neurodegenerative diseases^17–19^. Organoids derived from the airway epithelium have gained prominence for studying diseases affecting respiratory tissues such as the lung^20^ and nasal^21^ epithelia. Comprising hollow spheres of cell layers, these organoids mimic phenotypic characteristics of their source epithelium. Typically generated by suspending cells in a basement membrane matrix hydrogel, such as Matrigel, these cells subsequently differentiate and self-assemble into spherically shaped, epithelial structures^22^. Given that typical respiratory epithelia lack embedded neuronal cell bodies, these model systems primarily comprise basal cells, ciliated cells and secretory cells. Notably, to the best of our knowledge, no examples of OE organoids containing a neuronal cell population embedded within the epithelium have been reported to date.

In this study, we present a new technique, adapted from existing airway organoid methodologies, for generating olfactory organoids from mouse OE. These organoids exhibit similar shape and structural characteristics to airway organoids but also harbor neuronal cells representing OSNs, alongside other olfactory cell types such as sustentacular and secretory cells. Further, we demonstrate the expression of α-synuclein in these organoids, as an example of their value to Parkinson disease research. In addition, we demonstrate the model’s platform to study gene-environment interactions by inoculating murine organoids of OE with a neurotropic RNA virus.

## Results

### Dissection of mouse olfactory epithelium

Isolation of OE from 8-12 week-old C57BL/6J mice was conducted in accordance with protocols approved by the University of Ottawa Research Ethics Board (Protocol Number: ACVS NSIe-3557). Anesthesia was induced via intraperitoneal injection of euthanyl (120 mg/kg), followed by decapitation to ensure humane euthanasia. The step-by-step procedure for accessing and harvesting murine OE is depicted in **Figure 2**.

**Figure 2:**
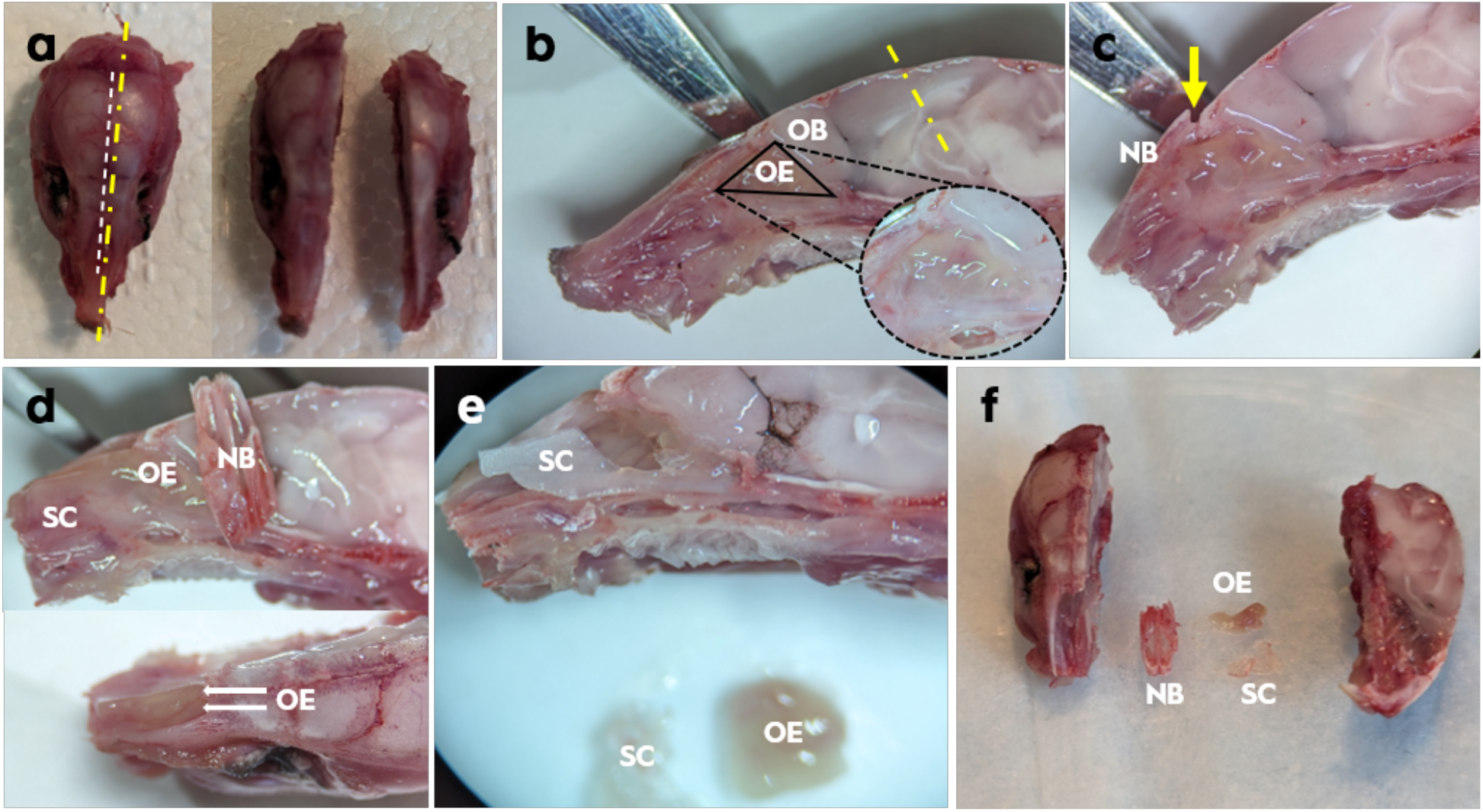
Dissection steps for collecting olfactory epithelia. **a)** Example of a mouse skull following removal of skin and mandible displaying the sagittal cut line (in yellow). **b)** Lateral view of the larger skull piece illustrating the triangularly shaped septal olfactory epithelium (OE) and the excision line for the nasal apex. **c)** Detachment of nasal bone (NB) after excision of the nasal apex. The yellow arrow indicates the dislodged cranial suture that allows removing the nasal bone while leaving the underlying OE intact. **d)** Lateral (upper panel) and superior (lower panel) appearances after NB removal depicting the septal cartilage (SC) and the surrounding OE. **e)** Lateral view following excision of OE and portion of SC. **f)** Comprehensive overview displaying dissected skull elements mentioned above.

The skin was peeled off the skull, and jaw and eyes were excised. The skull was then disinfected with 70% ethanol and carefully bisected along the sagittal plane, maintaining a 1 mm distance from the interfrontal suture. The larger segment of the skull, housing the nasal septum, was rinsed with phosphate-buffered saline (PBS) and positioned under a dissection microscope, with the nasal septum oriented upwards.

The OE, distinguishable by its yellowish hue in contrast to the white appearance of respiratory epithelia, was readily discernible as a triangular patch under the dissection microscope. Subsequently, a ∼1 mm distal section of the nasal bone was excised, which made it easier to detach tissue layers from the skull at the proximal site. Nasal bones were carefully removed to release the septal cartilage and surrounding OE. The latter was collected in its entirety using sharp-tipped forceps and transferred into Hanks’ Balanced Salt Solution (HBSS) (Stemcell Technologies 37250) for further processing.

### Olfactory epithelium stem cell culture

Culture plate surfaces were covered with 100 µg/mL poly-d-lysine hydrobromide (Millipore P6407) solution the day before cell isolation, as well as before passaging, and kept at 37°C. The next day, the poly-d-lysine hydrobromide solution was aspirated and surfaces were rinsed twice with sterile deionized water. After drying at room temperature, surfaces were covered with a 10 mg/mL laminin (Millipore L2020) solution in PBS containing Mg^2+^ and Ca^2+^. The culture dishes were incubated at 37°C for at least two hours before seeding cells and were not allowed to dry at any point.

Isolation and expansion of stem cells from harvested OE were adapted from the report by Peterson et al^10^. PneumaCult-Ex Plus Medium (PCP) (Stemcell Technologies 05040) was prepared according to manufacturer protocols. Collagenase/Hyaluronidase 10x mixture (Stemcell Technologies 07912) was diluted to 1x in PCP medium and the harvested OE from each mouse was transferred to 2 mL of this mixture. Pooling the harvested OE from multiple mice also yielded similar results but the numbers here were given for a single mouse. The mixtures were incubated at 37°C for 10 minutes and vortexed briefly before being centrifuged at 80 x g for 1 minute and aspirating the supernatant. All centrifuge steps were carried out at room temperature. The retrieved tissue pellet was triturated in 1 mL TrypLE Express Enzyme (1X) (Gibco 12604013) 30 times using a P1000 pipette. The enzyme was diluted by adding 5 mL HBSS and the cells were recovered by centrifuging at 350 x g. The pellet was resuspended in a mixture of 1 mL 1U/mL dispase (Stemcell Technologies 07923) and 50 µL 1 mg/mL DNase 1 solution (Stemcell Technologies 07900) and triturated briefly, until no visible tissue pieces remain. The enzymes were diluted once again with 5 mL HBSS and passed through a 50 µm cell strainer (Sysmex 04-004-2327) before centrifuging at 350 x g to obtain the final cell pellet. The expansion medium consisted of the PCP medium prepared according to manufacturer instructions further supplemented with: 2 mM L-glutamine (Gibco 25030081), 1x B27 supplement minus vitamin A (Gibco 12587010), 1x N2 supplement (Gibco 17502048), 10 ng/mL human recombinant TGF alpha (Stemcell Technologies 78123), 1 µM A-83-01 (Stemcell Technologies 72024) and 1 µM DMH1 (Stemcell Technologies 73634). The cells were plated in expansion medium on a surface area of 1.9 cm^2^ per dissected mouse.

As reported previously by Peterson et. al.^10^ cell colonies emerged around 3 days as phenotypically heterogenous clusters of tightly packed cells. Expression of globose basal cell marker LGR5 and horizontal basal cell marker CK14 were observed, as shown in **Figure 3b**. After reaching about 80% confluence, the cells were sub-cultured with 1:3 to 1:4 ratios in dishes coated with poly-d-lysine and laminin. Cells were detached from plates by incubating in Accutase (Stemcell Technologies 07920) for 5 min followed by trituration to gently break cell-surface and cell-cell adhesions. Detachment of cells using longer incubation times or other enzyme products such as TrypLE or trypsin resulted in loss of viability. Cells were expanded for at least 3 passages before being used for organoid induction or frozen for later use and expansion. Expansion medium supplemented with 5% dimethyl sulfoxide was used for cryopreservation.

**Figure 3:**
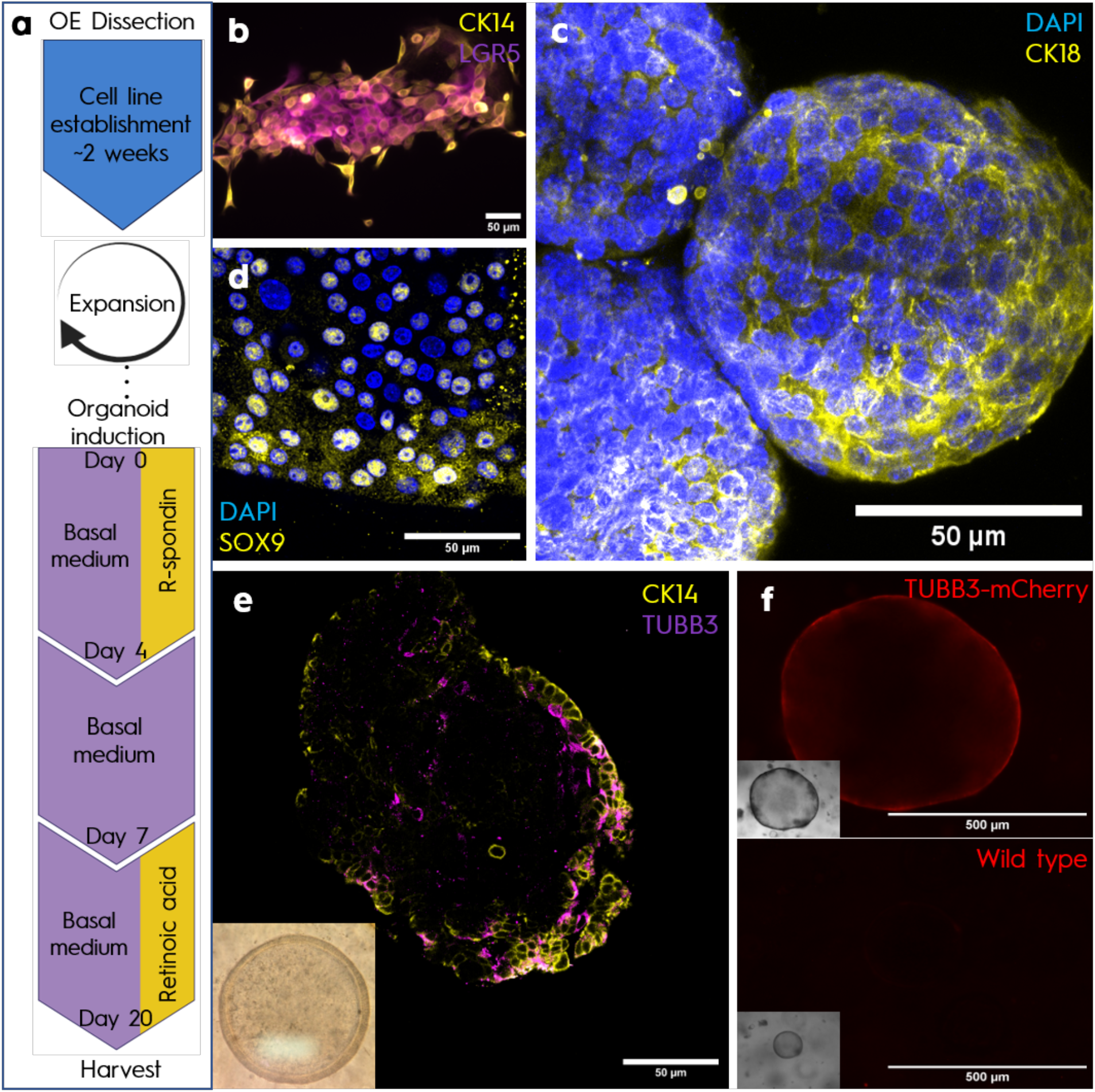
Development and characterization of olfactory epithelium (OE) organoids from mice. **a)** Summary of workflow for generating OE organoids; **b)** Early olfactory stem cell colony exhibiting heterogenous expression of CK14 and LGR5 proteins. **c)** OE organoids stained with anti-CK18 antibody demonstrating differentiation into a sustentacular phenotype. **d)** Organoid micrograph displaying the presence of SOX9+ secretory Bowman’s duct cells. **e)** Organoid stained with anti-CK14 and anti-TUBB3 revealing the organization of basal cells and presence of neurons. Inset shows a representative light microscopy image. **f)** Organoids derived from 8-week-old mice expressing transgenic TUBB3-mCherry, demonstrating fluorescence indicative of the presence of neurons compared to wild-type counterparts. Insets show brightfield photographs of organoids for the two distinct genotypes.

### Olfactory epithelium organoid culture

The protocol for organoid generation was adapted from Sachs et. al.^22^ wherein organoids are formed from cells suspended in droplets of a basement membrane matrix hydrogel. The basal medium formulation utilized in this protocol comprised a 1:1 mixture of Neurobasal A (Gibco 10888022) and advanced DMEM (Gibco 12491023) supplemented with 2 mM L-glutamine (Gibco 25030081), 1x B27 supplement minus vitamin A (Gibco 12587010), 1x N2 supplement (Gibco 17502048), 1 µM A-83-01 (Stemcell Technologies 72024), 1 µM DMH1 (Stemcell Technologies 73634), 3 µM CHIR 99021 (Stemcell Technologies 72054), 5 mM nicotinamide (Stemcell Technologies 07154) and 1.25 mM N-acetyl-L-cysteine (Sigma Aldrich A9165). During the initiation phase, basal medium was additionally supplemented with 500 ng/mL R-spondin-1 (Stemcell Technologies 78213).

To initiate organoid formation, olfactory stem cells were detached with Accutase, suspended in expansion medium and counted. The required volume from the cell suspension to obtain 4×10^4^ cells per well was transferred to a new tube and centrifuged. The cell pellet was resuspended in a 1:3 mixture of organoid initiation medium and matrix hydrogel precursor solution (Geltrex Reduced growth factor, Gibco A14132-02). 40 µL droplets of this cell suspension were dispensed into a 24-well plate that was pre-warmed at 37°C. The plate was quickly returned to 37°C and incubated for 20 min, until the matrix hydrogel set. Subsequently, the wells were gently flooded with 500 µL of initiation medium to prevent detachment of hydrogels from the well surface, and the plate was returned to the incubator (Day 0).

Self-assembly of organoids was observed as early as day 3. On day 4, the medium was refreshed for the first time with organoid basal medium without R-spondin-1. On day 7, after complete organoid formation, the medium was refreshed with organoid basal medium supplemented with 3 µM retinoic acid (Sigma Aldrich R2625). Of note, the addition of retinoic acid at an earlier stage inhibited organoid formation. FGF activation and MAPK p38 inhibition - typically included in airway organoids-were not found to be useful for olfactory organoids, and thus omitted. This medium exchange was repeated every 3 days until day 20 when the organoids were ready for harvest. The colony forming efficiency, defined as the ratio of emerged organoids to the starting cell number, at this time point was calculated to be 16.6% (± 3.4 SD).

Immunofluorescence microscopy at 20 DIV revealed the emergence of sustentacular cells as indicated by the presence of CK18 in **Figure 3c**. Expression of SOX9, a marker associated with secretory Bowman’s gland cells, confirmed the presence of secretory cells within the organoids (**Figure 3d**). Additionally, some cells positive for the neuronal cell marker TUBB3 were observed arising inward from the CK5+ basal cell layer as displayed in **Figure 3e**. To confirm the presence of neurons, organoids were generated from the cells of Tg-BAC-TUBB3-mCherry mice expressing the mCherry-reporter protein under the transcriptional control of the Tubb3 promoter. These mice were kindly provided by Dr. Minsheng Zhu. The mCherry signal was visible by live imaging, as shown in **Figure 3f**. Counterparts from wild-type mice monitored under the same live cell imaging conditions were not fluorescent, confirming the specificity of the mCherry signal, thus establishing the presence of neuronal cells embedded within organoids.

Expression of the Parkinson’s linked protein α-synuclein in the murine and human OE (**Figure 4a**) was previously shown by our group^23^ and others^24^. As an example of the potential of these organoids to study aspects of neurodegenerative diseases, we used immunostaining to confirm the presence of α-synuclein in olfactory organoids derived from wildtype mice, but not *Snca KO* mice (**Figure 4b,c**). We also confirmed the presence of α-synuclein in the conditioned media of olfactory organoids by ELISA^25^, whereas no signal was detected in conditioned media of olfactory organoids from *Snca* KO mice (data not shown).

**Figure 4:**
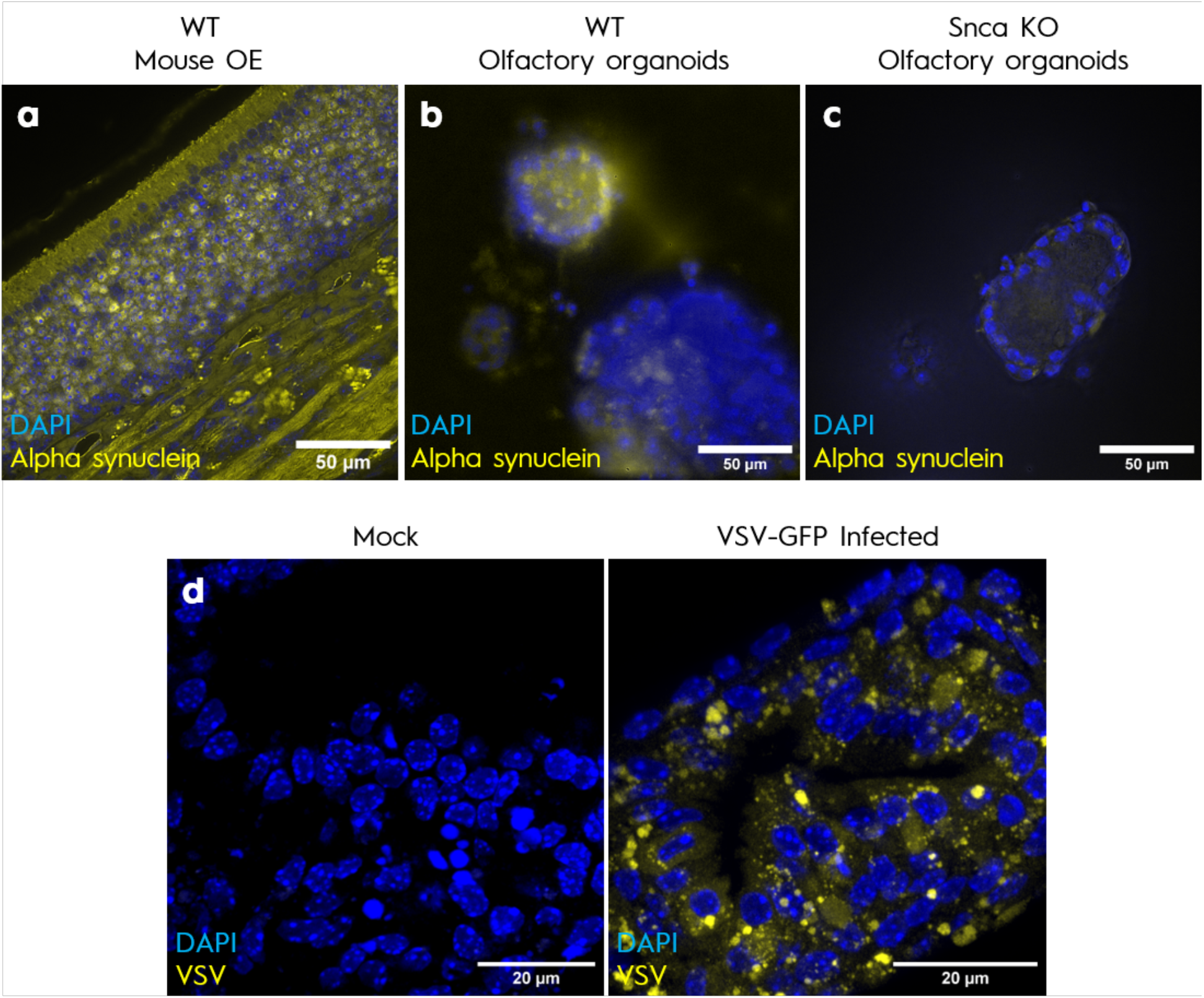
Generation of olfactory epithelium (OE) organoids for neurodegeneration-linked research purposes. Expression of Parkinson’s-linked α-synuclein visualized in: **a)** Intact OE and axons of cranial nerve-I, as shown in a sagitally dissected skull of a 6-month-old wild-type mouse; **b)** Wild-type olfactory organoids displaying expression of endogenous α-synuclein; **c)** Organoids generated in parallel from an α-synuclein-deficient knock-out (*Snca KO*) mouse as negative control of immunofluorescent signal shown in b. **d)** Olfactory organoids generated from wild-type mice treated with either medium alone (left panel; mock) or inoculated with 400,000 PFU of vesicular stomatitis virus (VSV-GFP, Indiana strain; right panel), as photographed by confocal microscopy 48 h after treatment induction.

Finally, to demonstrate the suitability of the organoids for studies involving environmental exposures such as viral infections, olfactory organoids were inoculated with a neurotropic ssRNA virus, i.e., GFP-tagged vesicular stomatitis virus (Indiana strain; VSV-GFP; a kind gift from Dr. Jean-Simon Diallo). As shown in **Figure 4d**, viral antigens were clearly expressed within the organoids after 2 days, in contrast to mock (PBS)-treated group, which demonstrated no viral protein expression.

### Collection, infection, staining and imaging of organoids

To harvest olfactory organoids, the medium was removed from wells, and 200 µL of dispase solution (5U/mL in HBSS, Stemcell Technologies 07913) were added to each well. The plates were then incubated at 37°C for 10 minutes, followed by gentle aspiration of the dispase solution without disturbing the hydrogels. Subsequently, 500 µL of PBS were added to each well and triturated using a P1000 pipette to release the organoids from the hydrogel. For this step and all subsequent transfers involving organoids, low-retention tips with the tips cut off were utilized.

For infection, organoids were first transferred to low-retention Eppendorf tubes and centrifuged at 100 x g for 4 minutes at room temperature. They were then placed into basal medium and infected with 300 µL of VSV-GFP at a concentration of 400,000 PFU per well, diluted in basal organoid medium. A control group (mock) received 300 µL of organoid medium. Organoids were subsequently moved to a 24-well plate and incubated for 48 hours. After this incubation period, the organoids were collected again in low-retention Eppendorf tubes and centrifuged at 100 x g at room temperature. The supernatant was discarded, and the organoids were fixed with a 4% formaldehyde solution (pH 7.2) at room temperature for 10 minutes before being transferred back to the 24-well plate. Between staining steps, the organoids were repeatedly transferred to low-retention Eppendorf tubes and centrifuged at 100 x g for 4 minutes at room temperature to facilitate solution aspiration by minimizing loss of isolated organoids.

For immunofluorescence staining, the organoids were initially incubated in staining buffer composed of 5% donkey serum (Biowest, S2170), 0.2% Triton X-100, and 0.1% Tween 20 for 1 hour. Next, primary antibody mixtures, prepared in staining buffer, were added at 4°C overnight. The sources of primary antibodies used in this study are listed in the Key Resource Table (**Supplemental Table 1)**.

Following overnight incubation, olfactory organoids were washed twice with PBS for 10 minutes each and then incubated with fluorescent secondary antibody mixtures for 2 hours at room temperature. They were then washed three times in PBS for 10 minutes each. Once the final wash was aspirated, 50 µL of ProLong Glass Antifade Mountant (Invitrogen P36980) was added to each tube, and organoids were mounted using uncoated #1.5 coverslips.

Images were acquired on a Zeiss LSM880 confocal commercial microscope with a fully motorized Axio Observer Z1 inverted stand, using a Zeiss 63x/1.4NA Plan-Apochromat DIC M27 oil immersion objective lens, Zeiss Laser launch light source and Zeiss AiryScan GaASP detectors. Images were acquired sequentially in linear scanning bi-directionally mode, using a voxel size of 0.25 µm x 0.25 µm x 0.82 µm and an area size of 135x135 µm in Zen Black 2.3 acquisition software (version 14.0.18.201), and were saved as CZI files.

## Discussion

In this study, we present an adaptation of existing airway organoid methodologies^22^ to generate olfactory organoids. Basal stem cells isolated from mouse olfactory epithelia were cultured and expanded routinely for up to three weeks, providing an abundant and pluripotent source of starting material for organoid formation. Detailed dissection methods are provided to aid researchers in efficiently harvesting mouse OE, addressing the limited knowledge and paucity of visual guidance available in the current literature.

Our organoid generation protocol involved modifying the medium formulation and administering specific reagents at different time points to induce differentiation and self-assembly of olfactory cell phenotypes, including of neuronal populations. This was made possible by adaptations that included a change in the base medium to a mixture of Neurobasal A and advanced DMEM to promote neuronal survival. Further, p38 MAPK blockers were omitted due to their inhibitory effect on neuronal differentiation^26^. Additionally, the activation of FGF signaling, commonly utilized in airway organoid cultures, was unnecessary as the starting cell population consisted of stem cells; Wnt signaling was activated via CHIR 99021 and an initial pulse of R-spondin-1. Notably, the inclusion of retinoic acid in the initial medium inhibited organoid formation but enhanced neuronal differentiation when added after the 7-day mark. Neuronal differentiation typically occurred around day 15, with all imaging analyses conducted at day 20.

Immunofluorescence micrographs confirmed the presence of basal cells, sustentacular cells, secretory cells, and neurons, each marked by corresponding cellular markers. Neuronal presence was further confirmed using organoids derived from transgenic mice expressing the TUBB3 promoter-driven mCherry reporter construct, which exhibited detectable mCherry fluorescence signals. Notably, OMP, associated with mature OSNs, was not yet detected in the olfactory organoids. OMP function is currently not fully understood, but it is linked to odor processing and glomerular integration of OSN axons^27^. Given its association with higher tissue hierarchy and established neural connectivity, which are attributes not (yet) recapitulated in our organoids that were followed until day 21 *in vitro*, the absence of OMP expression was anticipated. Notably, immature OSNs possess odor detection capabilities notwithstanding the absence of OMP expression^9^. Alpha-synuclein expression in the OE, previously demonstrated by our group^8^ and others^24,28^, was also detected in olfactory organoids, highlighting their relevance for studying Parkinson disease-related pathophysiology.

The relevance of studying the OE, particularly in the context of neurodegenerative diseases, is increasingly evident. Emerging evidence underscores the pivotal role of environmental factors in neurodegenerative diseases^29–31^, with the olfactory system serving as a vulnerable point of entry into the central nervous system, thereby potentially implicating it in disease initiation^32^. Current studies investigating the effects of environmental factors on humans primarily rely on epidemiological studies, which are limited in their ability to establish causal relationships or post-mortem studies which are restricted by the limited availability, tissue processing issues and non-acute exposure history. Organoid models hold promise for addressing these limitations. Notably, the olfactory organoid model developed in this study has proven to be suited for exposure studies. Once released from the basement membrane hydrogel, the organoids were readily infected by VSV added to the medium, indicating that this model is adaptable for exposure to a broad range of factors commonly employed in conventional *ex vivo* and *in vitro* studies. The method developed herein could be adapted to human cells obtained from biopsies or nasal swabs (provided that deeper layers containing proliferation-competent basal cells are sufficiently collected), which would enable the generation of human olfactory organoids.

Such mammalian model systems have the potential to facilitate the critical evaluation of a range of environmental factors associated with the epidemiology of human brain disorders. In conclusion, olfactory organoids provide an additional platform to study complex interactions between exposome and genome as they relate to the pathogenesis of select neurological disorders^33^.

## Data Availability

The protocols and lab materials used in this study are listed in a Key Resource Table alongside their persistent identifiers as **Supplemental Table 1**.

## Acknowledgements

This research was funded by Aligning Science Across Parkinson’s [Grant ID: ASAP-020625] through the Michael J. Fox Foundation for Parkinson’s Research (MJFF) and by the Parkinson Research Consortium Ottawa. We are grateful to the Department of Medicine at The Ottawa Hospital and the Uttra and Sam Bhargava Family for their ongoing support (to M.G.S.). The authors acknowledge the Cell Biology and Image Acquisition Core (RRID: SCR_021845) funded by the University of Ottawa, Ottawa, Natural Sciences and engineering Research Council of Canada, and the Canada Foundation for Innovation. Histology services were provided by the Louise Pelletier HCF (RRID: SCR_021737).

---

**Supplemental Table 1:**
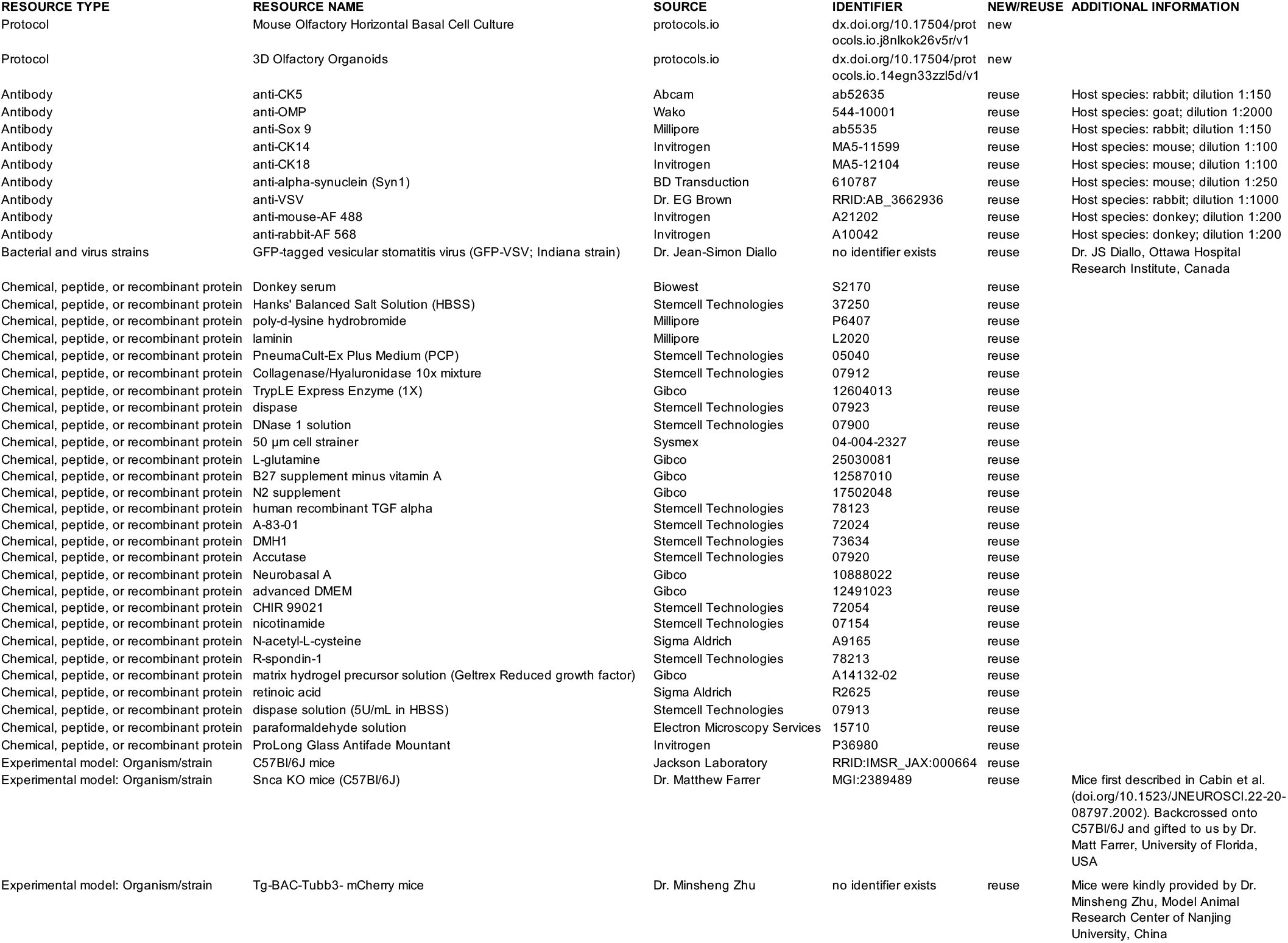
Key Resource Table for Ozgun et al., A Protocol for Neuralized Murine Olfactory Organoids.

