## Supplemental Table 1 for "A Protocol for Neuralized Murine Olfactory Organoids"

**Supplemental Table 1:** Key Resource Table for Ozgun et al., A Protocol for Neuralized Murine Olfactory Organoids

| RESOURCE TYPE | RESOURCE NAME | SOURCE | IDENTIFIER | NEW/REUSE | ADDITIONAL INFORMATION |
|---|---|---|---|---|---|
| Protocol | Mouse Olfactory Horizontal Basal Cell Culture | protocols.io | dx.doi.org/10.17504/protocols.io.j8nlkok26v5r/v1 | new | |
| Protocol | 3D Olfactory Organoids | protocols.io | dx.doi.org/10.17504/protocols.io.14egn33zzl5d/v1 | new | |
| Antibody | anti-CK5 | Abcam | ab52635 | reuse | Host species: rabbit; dilution 1:150 |
| Antibody | anti-OMP | Wako | 544-10001 | reuse | Host species: goat; dilution 1:2000 |
| Antibody | anti-Sox 9 | Millipore | ab5535 | reuse | Host species: rabbit; dilution 1:150 |
| Antibody | anti-CK14 | Invitrogen | MA5-11599 | reuse | Host species: mouse; dilution 1:100 |
| Antibody | anti-CK18 | Invitrogen | MA5-12104 | reuse | Host species: mouse; dilution 1:100 |
| Antibody | anti-alpha-synuclein (Syn1) | BD Transduction | 610787 | reuse | Host species: mouse; dilution 1:250 |
| Antibody | anti-VSV | Dr. EG Brown | RRID:AB_3662936 | reuse | Host species: rabbit; dilution 1:1000 |
| Antibody | anti-mouse-AF 488 | Invitrogen | A21202 | reuse | Host species: donkey; dilution 1:200 |
| Antibody | anti-rabbit-AF 568 | Invitrogen | A10042 | reuse | Host species: donkey; dilution 1:200 |
| Bacterial and virus strains | GFP-tagged vesicular stomatitis virus (GFP-VSV; Indiana strain) | Dr. Jean-Simon Diallo | no identifier exists | reuse | Dr. JS Diallo, Ottawa Hospital Research Institute, Canada |
| Chemical, peptide, or recombinant protein | Donkey serum | Biowest | S2170 | reuse | |
| Chemical, peptide, or recombinant protein | Hanks' Balanced Salt Solution (HBSS) | Stemcell Technologies | 37250 | reuse | |
| Chemical, peptide, or recombinant protein | poly-d-lysine hydrobromide | Millipore | P6407 | reuse | |
| Chemical, peptide, or recombinant protein | laminin | Millipore | L2020 | reuse | |
| Chemical, peptide, or recombinant protein | PneumaCult-Ex Plus Medium (PCP) | Stemcell Technologies | 05040 | reuse | |
| Chemical, peptide, or recombinant protein | Collagenase/Hyaluronidase 10x mixture | Stemcell Technologies | 07912 | reuse | |
| Chemical, peptide, or recombinant protein | TrypLE Express Enzyme (1X) | Gibco | 12604013 | reuse | |
| Chemical, peptide, or recombinant protein | dispase | Stemcell Technologies | 07923 | reuse | |
| Chemical, peptide, or recombinant protein | DNase 1 solution | Stemcell Technologies | 07900 | reuse | |
| Chemical, peptide, or recombinant protein | 50 µm cell strainer | Sysmex | 04-004-2327 | reuse | |
| Chemical, peptide, or recombinant protein | L-glutamine | Gibco | 25030081 | reuse | |
| Chemical, peptide, or recombinant protein | B27 supplement minus vitamin A | Gibco | 12587010 | reuse | |
| Chemical, peptide, or recombinant protein | N2 supplement | Gibco | 17502048 | reuse | |
| Chemical, peptide, or recombinant protein | human recombinant TGF alpha | Stemcell Technologies | 78123 | reuse | |
| Chemical, peptide, or recombinant protein | A-83-01 | Stemcell Technologies | 72024 | reuse | |
| Chemical, peptide, or recombinant protein | DMH1 | Stemcell Technologies | 73634 | reuse | |
| Chemical, peptide, or recombinant protein | Accutase | Stemcell Technologies | 07920 | reuse | |
| Chemical, peptide, or recombinant protein | Neurobasal A | Gibco | 10888022 | reuse | |
| Chemical, peptide, or recombinant protein | advanced DMEM | Gibco | 12491023 | reuse | |
| Chemical, peptide, or recombinant protein | CHIR 99021 | Stemcell Technologies | 72054 | reuse | |
| Chemical, peptide, or recombinant protein | nicotinamide | Stemcell Technologies | 07154 | reuse | |
| Chemical, peptide, or recombinant protein | N-acetyl-L-cysteine | Sigma Aldrich | A9165 | reuse | |
| Chemical, peptide, or recombinant protein | R-spondin-1 | Stemcell Technologies | 78213 | reuse | |
| Chemical, peptide, or recombinant protein | matrix hydrogel precursor solution (Geltrex Reduced growth factor) | Gibco | A14132-02 | reuse | |
| Chemical, peptide, or recombinant protein | retinoic acid | Sigma Aldrich | R2625 | reuse | |
| Chemical, peptide, or recombinant protein | dispase solution (5U/mL in HBSS) | Stemcell Technologies | 07913 | reuse | |
| Chemical, peptide, or recombinant protein | paraformaldehyde solution | Electron Microscopy Services | 15710 | reuse | |
| Chemical, peptide, or recombinant protein | ProLong Glass Antifade Mountant | Invitrogen | P36980 | reuse | |
| Experimental model: Organism/strain | C57Bl/6J mice | Jackson Laboratory | RRID:IMSR_JAX:000664 | reuse | |
| Experimental model: Organism/strain | Snca KO mice (C57Bl/6J) | Dr. Matthew Farrer | MGI:2389489 | reuse | Mice first described in Cabin et al. (doi.org/10.1523/JNEUROSCI.22-20-08797.2002). Backcrossed onto C57Bl/6J and gifted to us by Dr. Matt Farrer, University of Florida, USA |
| Experimental model: Organism/strain | Tg-BAC-Tubb3- mCherry mice | Dr. Minsheng Zhu | no identifier exists | reuse | Mice were kindly provided by Dr. Minsheng Zhu, Model Animal Research Center of Nanjing University, China |
